# Insecticidal roof barriers mounted on untreated bednets can be as effective against *Anopheles gambiae s.l.* as insecticidal bednets

**DOI:** 10.1101/2023.06.19.545532

**Authors:** Anthony J. Abbott, Agnes Matope, Jeff Jones, Vitaly Voloshin, Cathy Towers, David Towers, Philip J. McCall

## Abstract

Barrier bednets (BBnets), regular bednets with a vertical insecticidal panel to target mosquitoes above the bednet roof, where activity is highest, have the potential to improve existing Insecticidal Treated Bednets (ITNs), by reducing quantity of insecticide required per net, reducing the toxic risks to those using the net, thus increasing the range of insecticides to choose from. We evaluated performance of different BBnet variants based on the PermaNet 3 (*i*.*e*., P3 BBnets with pyrethroid and piperonyl butoxide (PBO) on the roof or barrier; pyrethroid alone on the side walls) in room-scale bioassays, simultaneously video-recorded to track mosquitoes. Experimental results showed the longitudinal P3 barrier (P3L) to be highly effective: P3+P3L were consistently though not significantly more effective than the reference P3 bednet while performance of Ut+P3L was comparable to the reference P3. Comparing contact duration at the treated sections of each variant, the Ut+P3L accumulated 1273 contacts with 1374 seconds duration, all on the barrier, greatly exceeding the 792 seconds duration, from 8049 contacts, accumulated across the entire surface of the PermaNet 3 reference bednet. The BBnet’s potential to augment existing bednets and enhance their performance is considered.

## INTRODUCTION

With its most effective control tool, the insecticide-treated bednet (ITNs), under serious threat from insecticide resistance, malaria control in Africa is at a critical stage. ITNs treated with pyrethroid insecticides were the primary method contributing to the impressive reductions in malaria incidence seen year on year prior to 2015. Since then, evidence of ITNs losing efficacy against resistant vector populations has become widespread while evidence of an association between pyrethroid resistance and re-emerging malaria is growing [1].

At the time of writing, there are 25 commercial ITN products listed with WHO prequalification, all of which use pyrethroids. A pyrethroid is the only active ingredient (a.i.) on sixteen of these nets, while on another seven, the added synergist piperonyl butoxide (PBO) disables the mosquito’s resistance mechanism, restoring pyrethroid efficacy. Only two ITNs deploy a second insecticide from a different insecticide class: a pyrrole chlorfenapyr, and juvenile hormone analogue, pyriproxyfen [2].

Clearly, PBO has extended the effective lifespan of pyrethroids as net treatments [3] and the absence of cross resistance between any major insecticide class and chlorfenapyr secures the immediate future for ITNs [4]. Historically, however, the arrival of resistance has been inevitable following the large-scale implementation of new net treatments, especially when genetically diverse and interconnected populations are being targeted [5]. If resistance to the new net treatments were to happen today, bednets would be left dependent on an insecticide class against which the target populations are already highly resistant [6], with poor prospects for expanding the range of potential treatments. Expansion of a list of available net treatments is essential if we are to ensure a future for ITNs in malaria vector control. However, the range of possible treatments remains constrained by the need to minimize risks to the sleeper, while the higher cost of new insecticides could make some treatments economically unviable. There are no alternative vector control options that can deliver levels of protection comparable to ITNs, both for those in the community that use nets routinely and those without nets. Failure to ensure their future would be disastrous for Africa.

In response to this challenge, bednets with barriers or ‘panels’ of insecticide treated netting on the ‘roof’ were designed to both improve the performance of a deltamethrin-treated bednet against pyrethroid-resistant mosquitoes and to permit the safe deployment of active ingredients [7]. Upright roof barriers intercept mosquitoes above the bednet roof, where *An. gambiae s*.*l*. are most active [8] and where the insecticide-treatment cannot contact the sleeper. The possibility that deploying a more effective insecticide on the barrier could result in a net capable of performances that are equal to those of a standard ITN is an attractive prospect. Such a BBNet could potentially increase the range of candidate insecticides, by making newer expensive insecticides affordable, allowing higher concentrations of the a.i. or permitting the use of some insecticides thar would be considered unsafe or unsuitable for direct skin contact if the sides and roof of the bed net were treated. In addition to increasing treatment choice, BBnets would require less insecticide per net, which translates as safer cheaper nets, less toxic waste and fewer impacts on non-target fauna.

Following the initial demonstration of barrier bednet efficacy against a wild pyrethroid resistant population in the field [7] we report here on laboratory tests investigating the potential of different roof barrier designs of PBO-treated netting. We used the same infra-red video tracking system as earlier studies [8,9] to video-record the interactions between *An. gambiae s*.*l*. from a laboratory colony as they flew freely in a large climate-controlled room, responding to a human volunteer host within different BBnet variants. This allowed considerable control over the experimental setup without interfering with the spatial behavior of the mosquitoes as it interacted with the ITN while responding to the host.

## RESULTS

The study started in late 2019 in UK and was forced to stop as the first Covid lockdown measures were implemented in March 2020, necessitating an abrupt curtailment in the number of repeat tests performed and the range of BBnet variants tested. The resistant Tiassale strain was tested with all BBnet variants, but the susceptible Kisumu strain was tested only with transverse BBNet types.

BBnet variant names are based on their composition: Bednet treatment (<u>U</u>n<u>t</u>reated or <u>P</u>ermanet <u>3</u>) + barrier treatment (Ut or P3) and shape (<u>t</u>ransverse ‘T’ or longitudinal ‘L’); *e*.*g*., Ut+P3L is an untreated bednet with a longitudinal barrier of Permanet 3 (P3) netting (Figs 1a, 2a).

**Figure 1.**
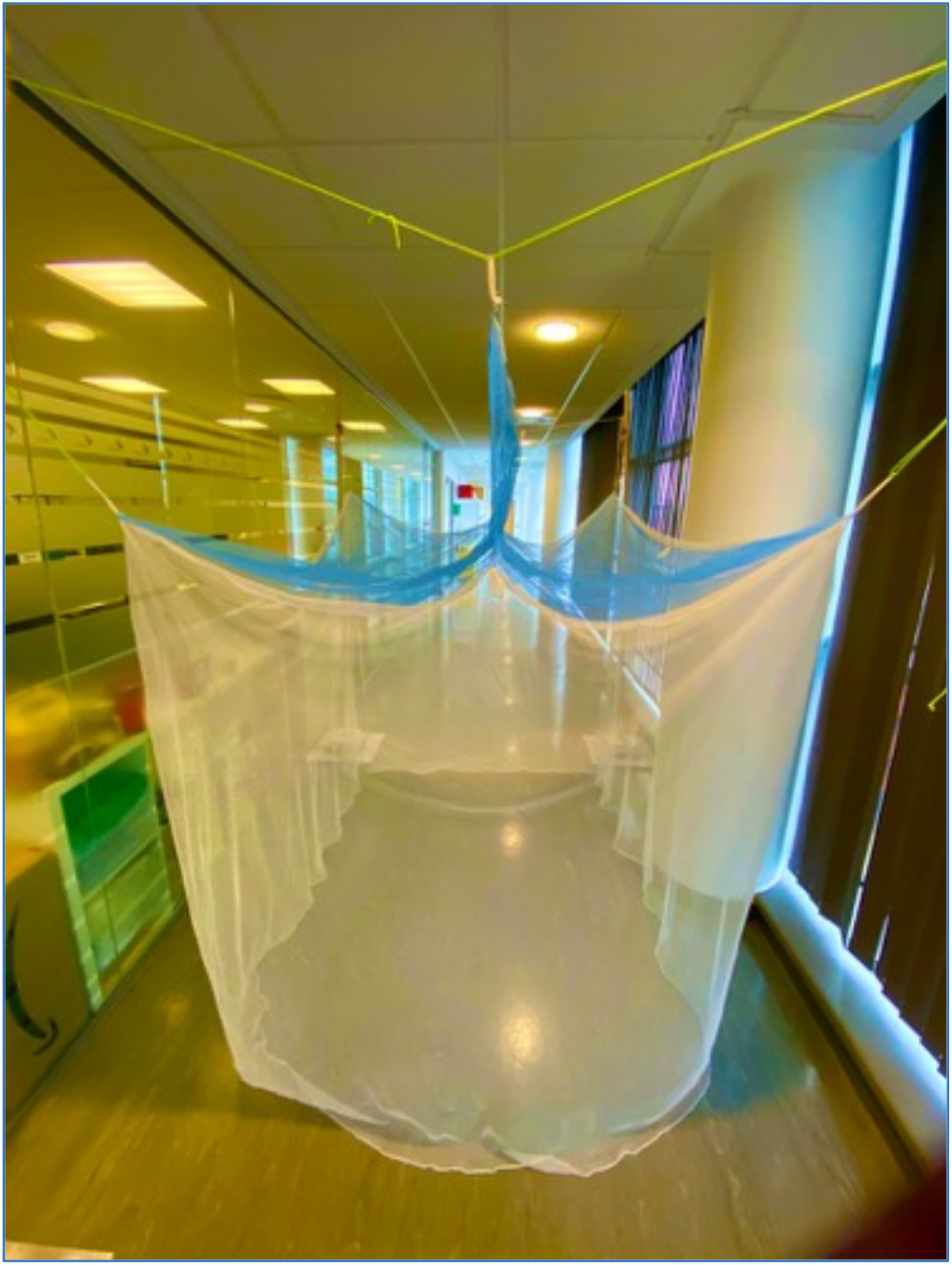
Example of a longitudinal Permanet 3 polyethylene barrier bednet. The blue netting on the barrier and roof is 100 denier polyethylene net with deltamethrin incorporated at 120 mg/ m^2^ and PBO at 750 mg/m^2^. The white netting on the sides is 75 denier polyethylene with deltamethrin incorporated at 84 mg/ m^2^.

Following completion of the assays, susceptibility to deltamethrin and permethrin of both strains was tested by 1hour exposure in WHO tube tests in May 2020 to determine their level of pyrethroid susceptibility Kisumu was fully susceptible to both, with 100% mortality recorded in all tests. However, Tiassale strain, originally selected for its high level of resistance to pyrethroids was only partially resistant, with mortality rates recorded post exposure of 13% and 8% to deltamethrin and permethrin, respectively.

### Performance of Transverse P3-BBnets against mosquitoes fully susceptible to pyrethroids

The mortality and knockdown rates of tests with susceptible mosquitoes at nets with transverse barriers are summarized in Fig 2. The unmodified P3 net killed 100% of mosquitoes in all but one test repeat, where it killed 97%. All barrier bednet combinations with at least one P3-treated surface knocked down 91-100% of susceptible mosquitoes within 1hr and killed 92-100% within 24hr (Fig 2b, C). Comparing the knockdown and mortality rates of the three BBNets with at least one P3 section, all the exposed Kisumu mosquitoes died within 24 hours post-exposure except for a few in the Ut + P3T. Although not statistically significant, the performance of the untreated bednet with a transverse P3 barrier (Ut + P3T) was comparable to a standard treated Permanet 3 ITN, indicating that the barrier’s position, above the net and directly over the human sleeper’s torso, is well placed to target *Anopheles sp* mosquitoes as they attempt to reach the human volunteer inside.

**Figure 2.**
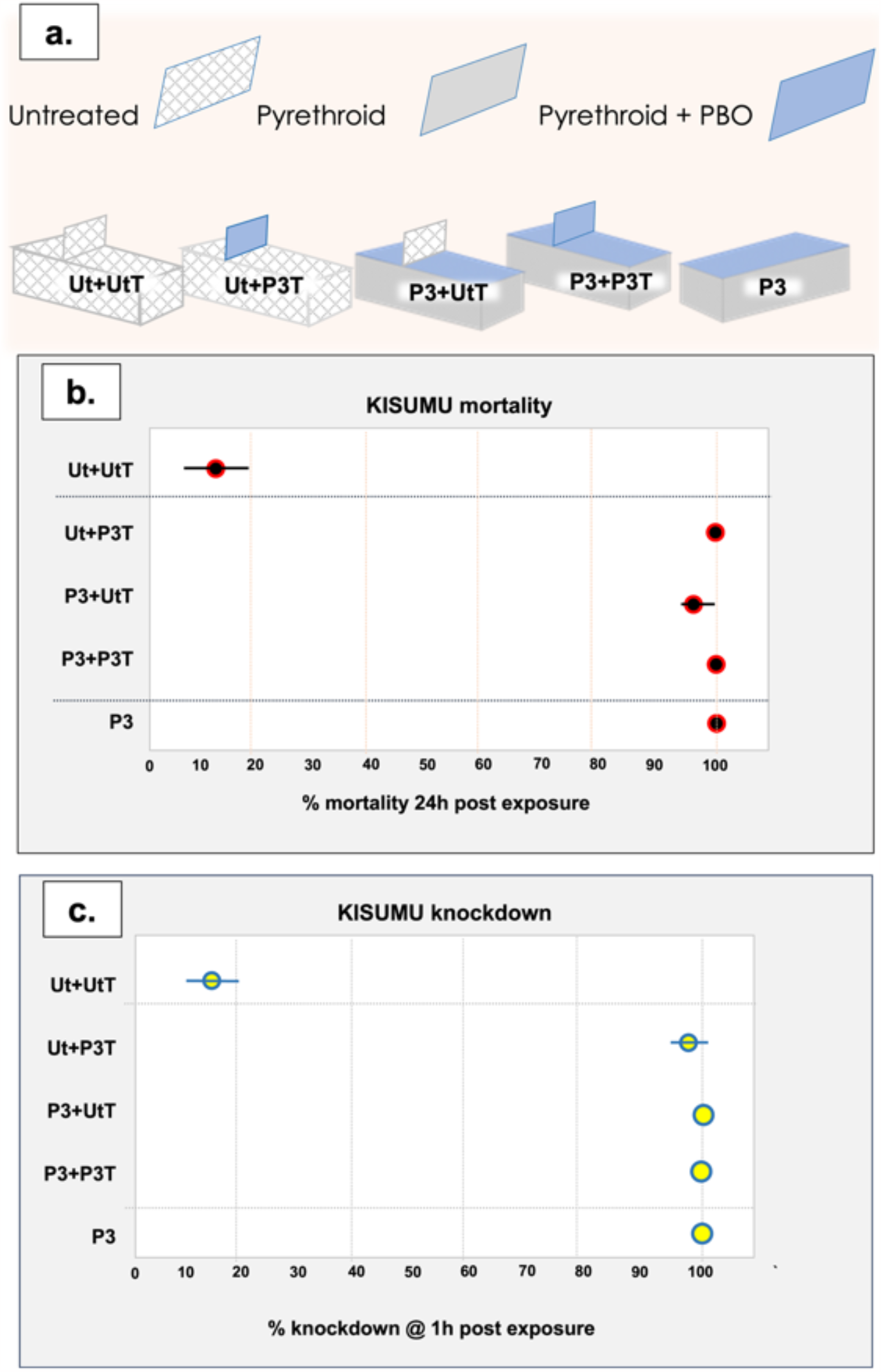
Mortality and knockdown rates of pyrethroid susceptible *Anopheles gambiae* Kisumu strain at transverse barrier bednet variants. **A)** schematic illustrating the composition of each of the BBnets tested; **B)** Mortality and **C)** knockdown rates following the room-scale tests, as determined from videos recorded during tests and final counts at the termination of the 120 min assay (mean and CI; n = 6 repeat tests /treatment).

### Performance of the BBnet variants against partially pyrethroid resistant mosquitoes

The mortality and knockdown rates of tests with Tiassale mosquitoes at transverse and longitudinal BBnets are summarized in Fig 3.

**Figure 3.**
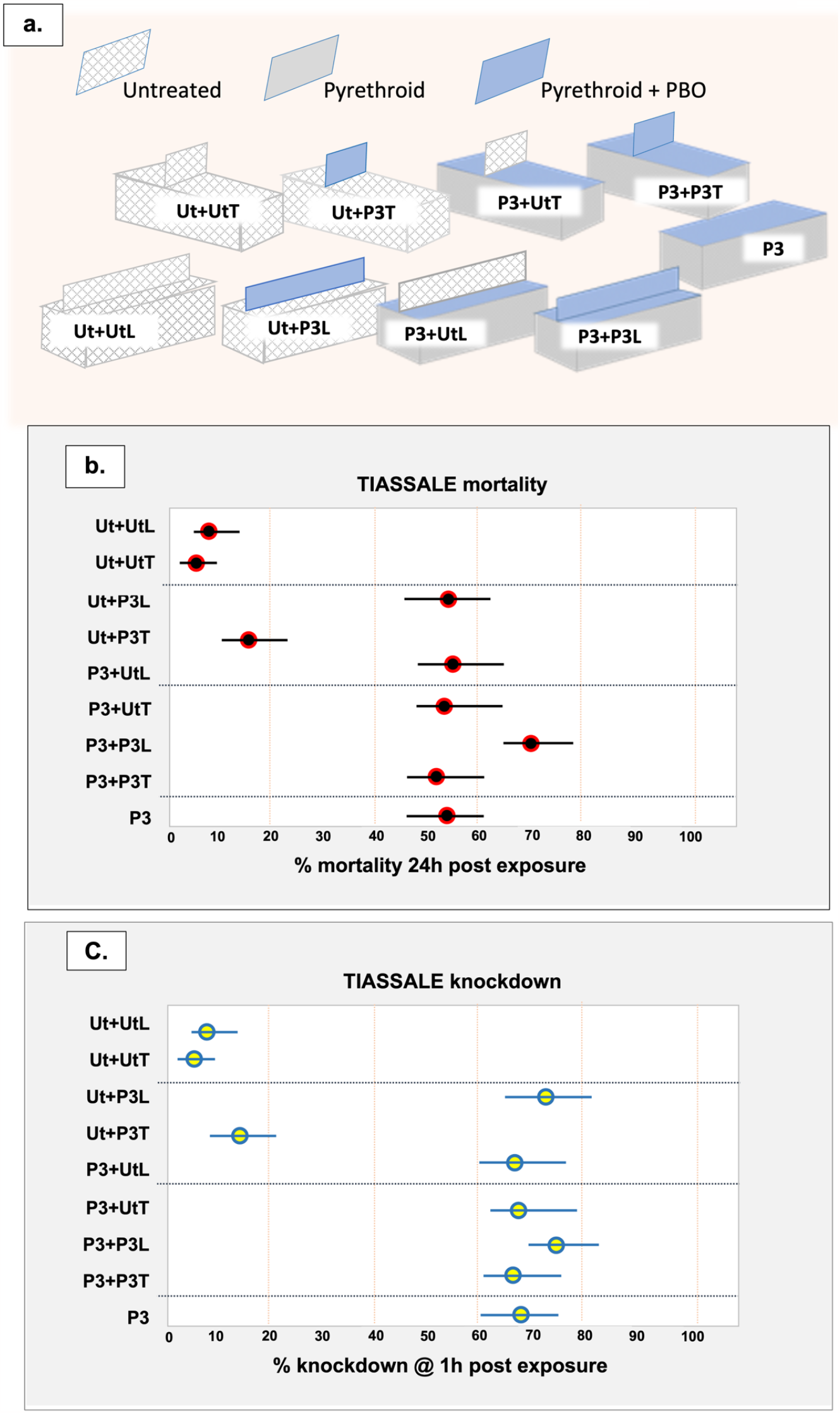
Mortality and knockdown rates of pyrethroid resistant *Anopheles gambiae* Tiassale strain at transverse and longitudinal barrier bednet variants. **A)** schematic of the composition of the BBnet variants tested. 1. **B)** Mortality and **C)** knockdown rates following the room-scale tests, as observed in videos recorded during the 120 min tests and final counts at the termination of the 120 min assay (mean and CI; n = 6 repeat tests /treatment).

Five BBnet variants achieved mortality rates greater than 50%, similar to the unaltered basic P3 reference (Fig 3b,c). The lowest performance was Ut+P3T, an untreated bednet with a shorter transverse P3 barrier. Tiassale mosquitoes were twice as likely to die when exposed to P3+P3L and 63% less likely to die when exposed to Ut+P3T compared to P3 (P3+P3L: OR = 2.09; 95% CI: 1.21, 3.64; *p* = 0.0086, and Ut+P3T: OR = 0.37; 95% CI: 0.20, 0.67; *p* = 0.0013). However, no significant differences between P3 and the other nets were found (Table S1).

Four of the five variants with efficacy comparable to the P3 (knockdown > 70%; mortality >50`%) comprised a barrier mounted on a complete P3 base. The fifth variant, Ut+P3L, was an exception, as it achieved mortality rates comparable to the other four variants and the P3 reference although its bednet base was untreated. The P3L barrier on a P3 bednet base also performed well, but although it exceeded mortality rates of the P3 in all assay repeats, the difference was not significant (wide confidence intervals OR = 0.93; 95% CI 0.57, 1.51; *p* = 0.7634).

#### Contact rate and duration of contact

The mean number of contacts (and CI) made and the duration of that contact for each BBnet variant are given in Figure 4, which shows the total number and duration of contacts on any part of each BBNet (fig 4a.) on the treated sections only (fig 4b.) and per cm^2^ of treated net (fig 4c). The duration of contact with the total net surface (all untreated and treated net surfaces, including barrier; fig 4a) during exposure was longer in all net variants compared to P3. However, these differences were significant only with P3+P3L, P3+UtL, Ut+P3L and Ut+P3T (borderline).

**Figure 4.**
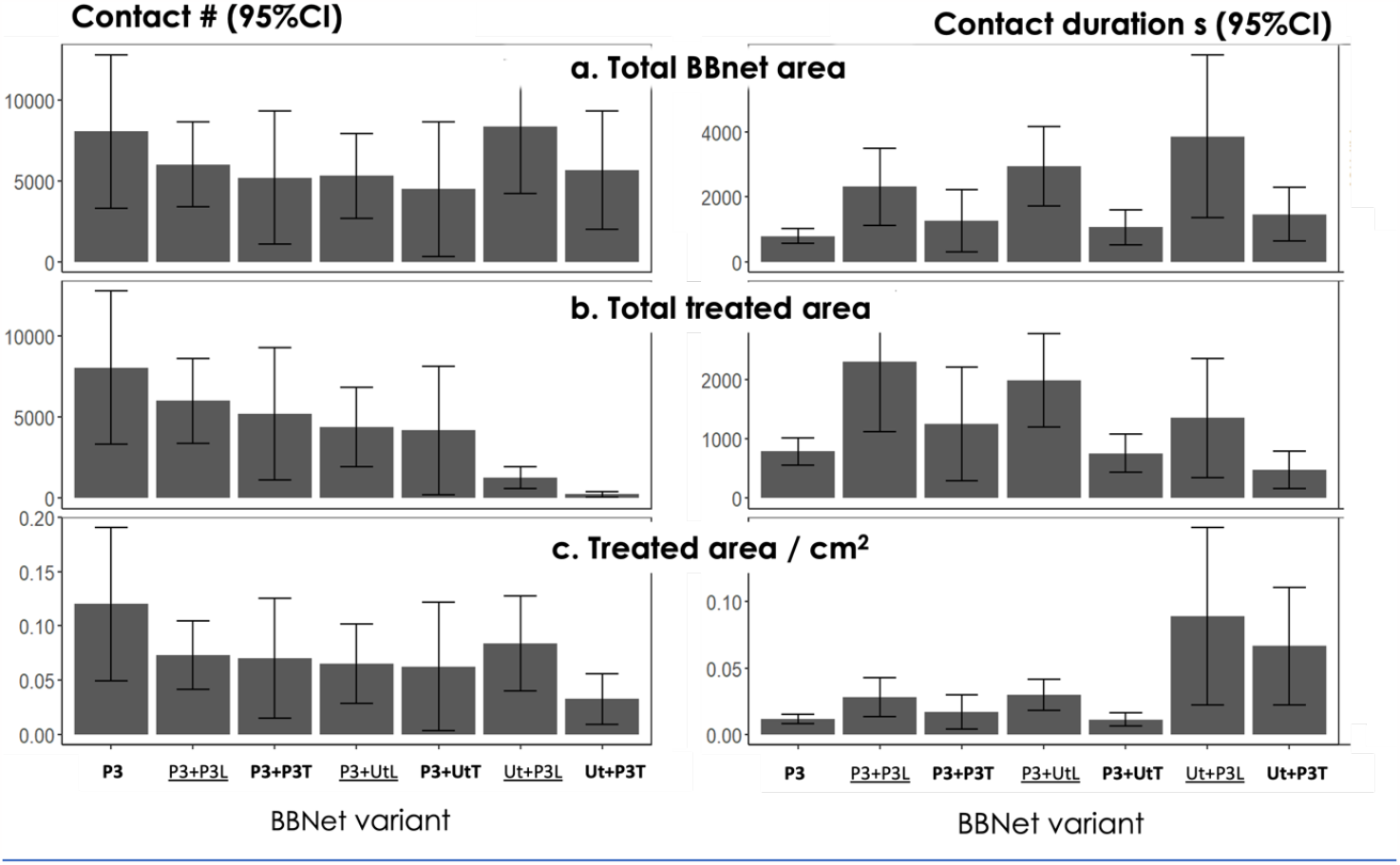
mean total numbers and durations of contact with the test netting during the 120 min assay for (a) all netting on the BBnet (b) all treated net surface area, (c) treated netting expressed per cm^2^.

**Figure 5.**
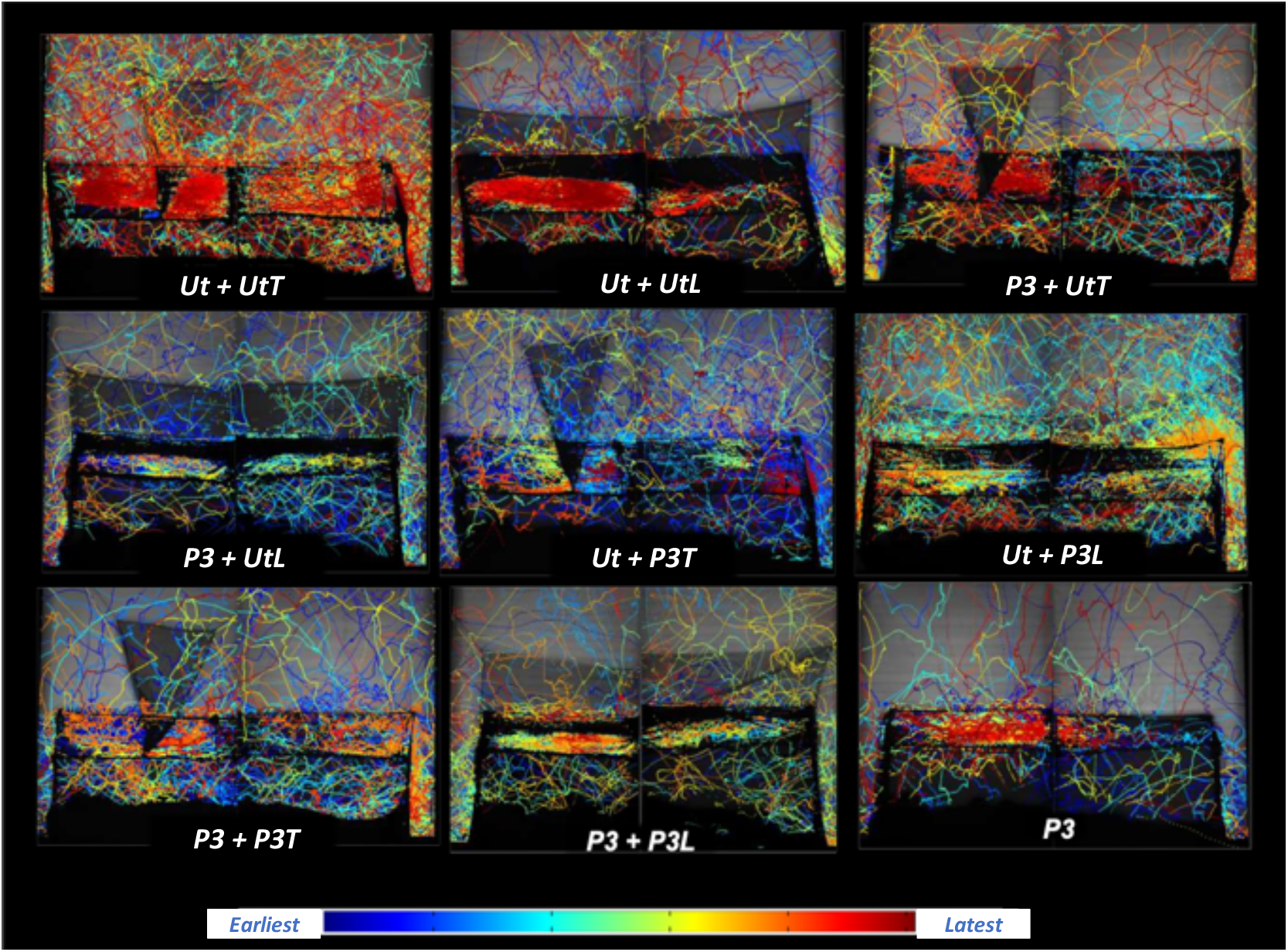
Examples of a full 120-minute composite images showing all flight tracks of *Anopheles gambiae* Tiassale strain at each of the BBnet variants. In all images, the sleeper is lying on their back, with their head at the left. Each colored track is the path of a single mosquito flight event, (25 mosquitoes released simultaneously) in all tests and color-coded according to time of appearance as shown in the key: blue tracks at the start through to red at the end of the 120-minute test.

The duration of contact with the treated net surfaces was significantly longer (at least 1465.66 secs) for Ut+P3L, P3+UtL, and P3+P3L compared to P3 net alone (Table S1, Ut + P3L: mean difference = 1465.99; 95% CI: 533.44, 2398.54; p = 0.0031, P3+UtL: mean difference = 1691.48; 95% CI: 885.42, 2497.54; p <0.0010, and P3 + P3L: mean difference = 1794.62; 95% CI: 1028.98, 2560.26; p <0.0010). Although we still do not know the precise minimum contact time required for a polyester ITN to deliver a lethal dose of an insecticide, increased contact duration is associated with higher kill rates [8].

Comparing contact duration at the treated sections of each bednet (fig 4b), the longitudinal P3 barrier on an untreated net accumulated 1374 seconds duration from 1273 contacts with, exceeding the 792 seconds duration at the Permanet 3 accumulated from 8049 contacts at the entire net surface.

## DISCUSSION

The results of these tests, although not fully conclusive, demonstrate the potential of simple roof mounted bednet barriers for malaria vector control. The bioassays clearly demonstrate that treated longitudinal barriers are likely to greatly improve the performance of the bednet beneath them. This ability could extend as far as ‘converting’ an untreated net into an effective ITN or ‘restoring’ an intact aged net simply by the addition of a treated barrier.

The key to the barriers’ ability to kill so efficiently is almost certainly their position on the roof: Here they are located directly above the supine human inside the bednet, where activity by incoming mosquitoes is greatest [8] maximizing the chance of encountering and interrupting flight paths of the majority of *Anopheles gambiae* visiting the net. This is illustrated by a comparison of the levels of contact at a longitudinal barrier with a total area of insecticidal netting of 7,600cm^2^ and a Permanet 3 bednet which has 8 times greater area of treated net, 62,200 cm^2.^ A comparison of the total duration of contact at the treated sections on a Ut + P3L BBnet and the reference P3 bednet (fig 4b), shows that despite the disparity in insecticidal surface area the longitudinal P3 barrier accumulated 1374 seconds duration from 1273 contacts, greatly exceeding the 792 seconds duration, from 8049 contacts, accumulated by all the treated net surface area at the Permanet 3. This was equivalent to contacts rates at the L barrier with a total duration of 0.9 s/cm^2^ compared with the P3 ITN which was 0.11s/cm^2^ (Fig 4c). The duration rates are remarkably similar, despite the difference in area.

The ITN has 8 times greater surface area loaded with insecticide, most of which does not contribute to its impact (fig 4). Moreover, mosquitoes orienting towards the bednet roof probably arrive first at the barrier / roof and may already have picked up a lethal dose of insecticide before they visit any other part of the net. This begs the question of whether an effective barrier on an untreated bednet base could kill mosquitoes as well as a regular treated bednet or put another way, could an untreated net reproduce the killing effect of a standard ITN simply by the addition of a treated barrier? These results certainly suggest it is possible, with the most appropriate insecticide treatment. To date, the BBnet has been tested mainly with insecticides that would be permissible on a standard bednet, except for the use of fenitrothion in Burkina Faso study and when the BBnet performed very well, increasing lethality without compromising personal protection [7].

The pattern of movement around a bednet where most mosquito activity occurs on or above the roof, is a response to olfactory and thermal cues rising from the host below, Sutcliffe & Yin [10] reported that the pattern disappeared when a breeze was blown across the host, dissipating any rising attractants, and eliminating the focus of activity on the net roof. This is true and it is widely known that sleeping with a steady low continuous flow of air from a reliable electric fan prevents mosquitoes from landing to the extent that it is possible to sleep comfortably without any bednet. However, at present few communities in endemic malaria zones are likely to have access to affordable power needed to run an electric fan all night every night, while security concerns are likely to continue to overrule any desire for comfort when deciding to open a window. Hence, until such improvements are widespread, this is unlikely to be a factor affecting the performance of BBNets.

The poor performance of the transverse barriers against the partially resistant mosquitoes was unexpected. In an earlier field study, we found that a transverse barrier treated with the organophosphate fenitrothion was highly effective against a wild population of pyrethroid-resistant *Anopheles gambiae s*.*l*., in Burkina Faso increasing killing approximately 34% more than a standard bednet [7]. Here, it performed well against susceptible mosquitoes (Fig. 2) but did not improve the performance of bednets against partially resistant mosquitoes compared to the negative controls. Differences in the insecticides used could explain this at least in part. In the field study where the transverse barriers were effective, they carried a highly effective insecticide, either deltamethrin against susceptible mosquitoes, or an organophosphate, fenitrothion against the wild highly resistant vector population in Burkina Faso [7]. In both cases, the insecticide’s effect was rapid following brief contact with a treated net. In studies where the transverse barriers performed poorly, as with Tiassale in the present study, impact depended on the synergist piperonyl butoxide (PBO) disabling the resistance mechanism before the insecticide could function. Although the minimum contact times needed to pick up a lethal dose for each of these treatments are not known, it is conceivable that the faster acting insecticides were more suited to delivery on a barrier, where the threshold for a lethal level of contact is brief (<50s)[11].

The loss of the highly resistant Tiassale colony when it was too late to repeat the experiments of was most unfortunate, given the importance of demonstrating an impact with resistant mosquitoes. However, as mentioned above, the Burkina Faso study where fenitrothion-loaded T-barriers were effective against highly resistant wild /natural vector population serves to demonstrate that BB-nets can be effective against highly resistant vectors [7].

With careful management, ITNs can remain the primary means of reducing malaria transmission in Africa for many more years. Standard bednet shapes can safely deliver only a very limited range of insecticides, seriously limiting options for management of insecticide resistance. Simplicity has been integral to ITN’s success, and nets do not need much sophistication to improve them. They employ the sleeper’s attractants to lure potentially infectious mosquitoes to the net surface where they are killed on contact while the sleeper is protected from bites behind a protective insecticidal screen. Requiring only minimal change to the basic design, a roof barrier exploits this further. Delivering insecticide on the roof alone also reduces the risk of the occupant’s skin becoming irritated by insecticide picked up during entry and exit from the protective net, further enhancing its advantages. With an estimated 79% of malaria cases transmitted when people are in bed [12], the value of a bed net is clear.

Given the importance of the net roof, and the region above it, for the lethality of any ITN, it would make sense if attempts to improve killing efficiency focused on this region. The low levels of mosquito activity at the vertical sides and ends of the bednet under normal conditions [8] suggest that the best we can do at these net locations, where most physical damage occurs, is to improve the quality of the material used in manufacturing the net to make stronger more durable nets.

Better bednets should mean a lot more than simply bednets with active ingredients that are effective against the target vector population, as this is surely the least any bednet should be. It is often a shock when the delicate fibers comprising the flimsy mesh that an ITN is made from are first seen. Sufficient knowledge of the entomological mode of action of ITNs already exists to enable us to identify the location of regions of the net to where mosquitoes are attracted, areas where the insecticide is best delivered to mosquitoes, vulnerable areas that require tougher net fibre, and areas rarely/never visited by mosquitoes. Can we not use this knowledge to design better bednets, ITNs that are optimised to prevent malaria transmission in rural Africa, rather than simply the cheapest product that ticks a few necessary boxes on a WHO form?

## MATERIALS & METHODS

### Mosquitoes and bioassays

All colonies were maintained, and bioassays performed in a climate-controlled unit at LSTM (27 ± 2°C, relative humidity 70 ± 10%) using 2-7-day old unfed adult females from the *Anopheles gambiae* s.l colonized strains, Kisumu (susceptible to all insecticides), and Tiassalé (part resistant to pyrethroids). Mosquitoes were deprived of sugar and water for 24h and 4h respectively before the experiment and transferred in a 300ml paper cup to the bioassay room to acclimatize for 1h prior to tests. Tests started 1-3 hours after the start of the scotophase with the release of test mosquitoes, were 2hrs in duration, after which time the room was searched thoroughly to count and collect live, dead or knocked down mosquitoes. All live mosquitoes were aspirated into a cup and provided with 10% sugar water in a separate room where mortality was recorded at 1h and 24h after the end of the experiment.

Volunteers, recruited from within LSTM, lay without shoes but fully clothed and uncovered within the bednet. They were requested to eschew strongly aromatic or spiced foods and scented personal hygiene products for 24h prior to each experiment. Since the tests were also video recorded for tracking, volunteers were requested to remain as still as possible for the duration of the experiment.

### BBnet variants

In all tests, rectangular bednets measuring 1.9 m × 0.8 m × ∼1.0 m tall were used as the standard bednet. Treated nets were Permanet 3.0 (Side walls of 75 denier polyester, deltamethrin 2.1g/kg±25% ; roof of 100 denier polyethylene, deltamethrin 4.0g/kg ± 25% and PBO 25g/kg ± 25%; Vestergaard, Lausanne). Barriers comprised sections cut to size from the roof of a Permanet 3. Untreated polyester nets were used as untreated controls, and matched Permanet 3 nets as closely as possible fiber thickness and hole size. Before being used in experiments, untreated nets were first tested to confirm the absence of any insecticidal effect, and all nets were hung for 4 weeks before use to allow evaporation of any potentially repellent or attractant volatile odors.

Two types of barrier were investigated, a transverse (from elbow to elbow of the supine sleeper beneath) and a longitudinal barrier (from head to toe), designs derived from earlier bioassay in the field or from mathematical models comparing their potential [7]. Transverse barriers were positioned off-center on the roof above the sleeper’s stomach or torso at the 30:70 division of the bednet’s length, where mosquito activity is known to be greatest [8].

To facilitate image capture on the top of the bednet, the net roof was tilted on its long axis when facing the cameras to ensure all mosquito activity on the roof was visible [8]. Hence, the height of the roof above the mattress was 0.80m at the front and 1m at the rear (when viewed/recorded from the front). To ensure the top edge was horizontal when mounted on this roof, the transverse barrier was 40cm tall at the rear and 60cm at the front of the net. The longitudinal barrier was 0.4m higher than the roof and ran the length of the bednet in the center of the roof.

The basic structure of transverse and longitudinal was similar in all tests as illustrated in Fig. 1a and 2a. Longitudinal BBNet designs were each tested in 6 repeats with both susceptible and resistant strains, but the transverse barriers were tested in 5 repeat tests with the Kisumu susceptible strain only, the result of time limits as the first COVID-19 lockdown approached. In the absence of further funding, the assays were not continued post-pandemic.

### Tracking Protocols

All bioassays were recorded for subsequent analysis of bednet contact using the video tracking system installed in Liverpool. This comprised of two identical cameras (Ximea CB120RG-CM with Canon EF 14 mm f/2.8L II USM lenses) with a custom LED ring attached (12 × OSRAMTM SFH 4235 infrared LEDs (peak wavelength 850 nm)). These were aligned with a Fresnel lenses per camera mounted at 1.2 m distance from the camera lens, and a retroreflective screen (plywood board, with 3M Scotchlite High Gain 7610 tape) mounted behind the bednet, 2 m from the Fresnel lenses. Recordings were made at 50fps using StreamPix software (www.norpix.com) and files saved in a .seq file format.

Recordings were segmented, creating a text file containing the coordinates and identity of each track, and additional noise recorded (such as movement from the volunteer) was cleaned up using the bespoke ‘seqfile processing’ software. The cleaned position files were then analyzed using bespoke MatLab (mathworks) software which defined the net regions. Continuous tracking of individual mosquitoes was not possible since the whole room is not in view, so each flight track was analyzed from the point of entry until leaving the field of view. The area recorded was separated into different regions breaking up different parts of the bednet and the surrounding area, with the number and duration of net contacts (bednet treated/ untreated or barrier treated/ untreated) recorded for each.

## Statistical analyses

Descriptive statistics were generated using the number of observations, mean, and standard deviation (SD) for continuous variables, the number and percentage of observations for categorical variables.

The duration of contact and the number of contacts made with each net were analysed using Linear Mixed Models (LMM). Mixed models were employed to account for the correlation between measurements from the same sleeper. Net type (nine categories: P3, P3 + P3T, P3 + P3L, P3 + UtT, P3 + UtL, Ut + P3T, Ut + P3L, Ut + UtT, Ut + UtL), maximum number of mosquitoes in contact with the net, the position of the volunteer’s head (two categories: left and right) and the possible significant interactions between these factors were included as fixed effects, and volunteer as a random effect.

For 1h KD and 24h mortality, a logistic regression model with a logit link function was fitted. The models for the Kisumu and Tiassale strains were fitted separately. For the Kisumu strain, the penalised likelihood estimation was employed to account for the complete separation that was observed for some net groups in the study [13]. The fixed effects considered included net type, duration of contact with the net/treated surface (Tiassale strain model only), and their interaction.

Statistical analyses were performed using R Version 4.2.1 and RStudio [14,15]. All the models were fitted using the “lme4” package while net contrasts were explored using the “emmeans” package [16,17]. Model selection was implemented on the full models described above to obtain the most parsimonious model. Likelihood ratio tests under the restricted maximum likelihood estimation were employed to determine the statistical significance of the random effects. The fixed effects for the ‘best’ fit models were selected based on model fitness determined using the Akaike information criterion and the residuals based on the “DHARMa” package [18]. Data were plotted using packages “ggplot2” and “ggfortify” [19, 20].

All the net types were compared to P3 (as reference), and the Dunnett multiple test adjustment procedure was employed to control for the probability of making false positive findings. The mean differences and odds ratios (OR) together with their corresponding 95% confidence interval (Cl) were generated for the net comparisons from the LMMs and logistic regression models, respectively. All the statistical tests were conducted at 5% Significance level.

## Ethical considerations

Since the participants would be protected within intact ITNs and were neither required nor expected to be bitten during the tests, full ethical clearance was not required by LSTM Research Ethics Committee. However, written informed consent was provided by all volunteers before participation.

## Acknowledgements

We thank Melinda Hadi for many informative discussions on the reality of bednet manufacturing and we are grateful to Vestergaard Sárl (Lausanne Switzerland) for part funding the study. Vestergaard had no role in the design, execution, analysis or reporting of the study.

## Author contributions

A.A. performed the experiments, collated outputs, and contributed to statistical analysis and writing drafts; A.M. performed the statistical analyses and interpreted results with A.A. and PJM; J.J. contributed to the BBNET Design and interpreted results; D.T. and C.E.T. designed the video-tracking capture and analysis system, which V.V. supervised and maintained; P.J.M. designed the BBnet, planned the experiments and wrote the manuscript with the input of the other co-authors, all of whom read and approved the final version.

## Data availability statement

Data are available from P.J.M. to whom all correspondence and requests should be addressed.

## SUPPLEMENTARY INFORMATION

### Abbott et al Barrier Bednets for malaria vector control

**Figure S1.**
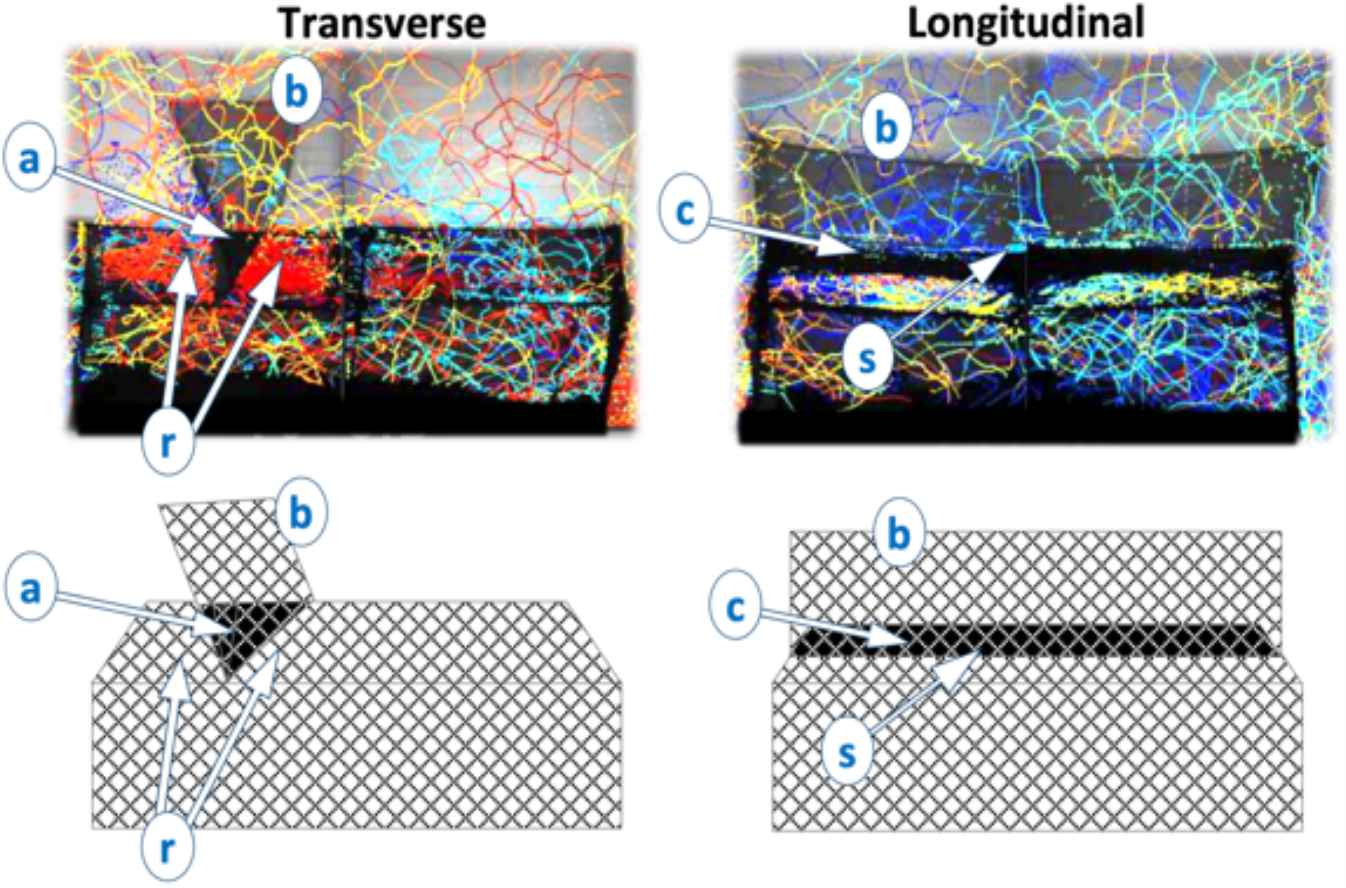
Areas where losses of multiple flight tracks required adjustment to compensate. Adjustments to tracking data to compensate for concealed tracks. The tracking system used here was the later refined system using a retro-reflective screen (RRS) to give more uniform and diffuse illumination (‘V2’; Voloshin *et* al, 2020). The original approach was back-lit with the light making a single pass through the measurement volume (‘A1’; Angarita-Jaimes *et al*., 2016). Using the RRS setup here with a BBNet we found that in certain sections, the position of a barrier added another layer of netting such that the line of sight of the camera passed through three layers of netting – and due to the retro-reflective imaging setup – these layers are double passed before the image is obtained at the camera. Such image areas gave few or no detected tracks (A and C in Fig S1). Each layer of net can be thought of as attenuating the optical signal and hence using RRS imaging with 3 net layers gives 6 traversals of net between the light source and the camera. For more conventional bednet setups when imaging through 2 layers of net (4 traversals), the RRS setup gave improved performance compared to back-lit imaging due to the diffuse nature and more uniform illumination produced. These benefits are lost when an additional bednet layer (the barrier) is added as the optical signal is reduced to give insufficient image contrast to segment the mosquito shadows from the already low optical signal. The back-lit imaging setup may have given better results for the BBNet case under study here, however, back-lit imaging requires 2 large area Fresnel lenses per camera and the lenses were already in use for studies in west Africa and committed to remain there for a lengthy period. To compensate for tracks lost during recording, track counts were adjusted as follow**s:** <u>Transverse barrier</u>: as mosquito activity in Section A was viewed through three layers of netting, much activity was concealed, and counts affected in areas equivalent to a mean of 10.5% of the area in the head end section of the roof (R) and 14.2% of the barrier (B). The final net contact counts for the barrier and the roof-head end were increased proportionately. <u>Longitudinal barrier</u>: here, Section C was obscured by 3 net layers in areas equivalent to mean rates of 42% of the roof (R), 29% the barrier (B) and the upper 50% of the sides: net contact counts were increased proportionately. Accuracy in adjustment was aided by the knowledge that distribution of tracks on bednets is equal on left and right sides of the host (Parker et al, 2015).

**Table S1.** 1h knockdown and 24h mortality results for all mosquitoes in tests with all BBnet variants. based on the logistic regression model with a logit link function. Where 95% confidence intervals and *p-values* were adjusted by Dunnett’s multiple comparison test, * = significant at 5% significance level, ^a^ = net reference P3, ^b^ = borderline significance, ^f^ = logistic regression based on the firth method/penalised likelihood and ref = reference.

| Term | Odds Ratio [95% Confidence Interval], p-value |  |  |  |  |  |
| --- | --- | --- | --- | --- | --- | --- |
|  | 1h KD |  | 24h mortality with all groups |  | Treated surfaces groups only (Tiassale) |  |
|  | Kisumu | Tiassale | Kisumu | Tiassale | 1h KD | 24h mortality |
| (Intercept) | 293.00<br>[42.98, 36877.75],<br><0.0010* | 2.13<br>[1.51, 3.03], <0.0010* | 293.00<br>[42.98, 36877.75],<br><0.0010* | 0.11<br>[0.05, 0.26], <0.0010* | 1.79<br>[1.22, 2.66], 0.0034* | 0.11<br>[0.05, 0.27],<br><0.0010* |
| Net: P3 + P3T<sup>a</sup> | 0.67<br>[0.00, 124.48], 0.8435 | 0.88<br>[0.53, 1.48], 0.6339 | 0.67<br>[0.00, 124.48], 0.8435 | 1.07<br>[0.64, 1.77], 0.8003 | 0.80<br>[0.48, 1.36], 0.4087 | 1.08<br>[0.65, 1.79], 0.7751 |
| Net: P3 + P3L<sup>a</sup> |  | 1.50<br>[0.90, 2.53], 0.1197 |  | 2.09<br>[1.21, 3.64], 0.0086* | 1.09<br>[0.60, 2.01], 0.7776 | 2.18<br>[1.22, 3.92], 0.0087* |
| Net: P3 + UtT<sup>a</sup> | 0.94<br>[0.01, 173.65], 0.9748 | 0.81<br>[0.48, 1.35], 0.4108 | 0.94<br>[0.01, 173.65], 0.9748 | 1.15<br>[0.70, 1.92], 0.5785 | 0.81<br>[0.48, 1.36], 0.4246 | 1.15<br>[0.69, 1.92], 0.5853 |
| Net: P3 + UtL<sup>a</sup> |  | 0.98<br>[0.60, 1.61], 0.9421 |  | 1.53<br>[0.91, 2.58], 0.1085 | 0.75<br>[0.43, 1.33], 0.3270 | 1.57<br>[0.91, 2.72], 0.1027 |
| Net: Ut + P3T<sup>a</sup> | 0.14<br>[0.00, 1.43], 0.1052 | 0.39<br>[0.24, 0.62], <0.0010* | 0.11<br>[0.00, 1.01], 0.0506<sup>b</sup> | 0.37<br>[0.20, 0.67], 0.0013* | 0.41<br>[0.25, 0.67], <0.0010* | 0.36<br>[0.20, 0.66], 0.0011* |
| Net: Ut + P3L<sup>a</sup> |  | 1.45<br>[0.87, 2.45], 0.1597 |  | 0.93<br>[0.57, 1.51], 0.7634 | 1.30<br>[0.77, 2.23], 0.3265 | 0.94<br>[0.58, 1.54], 0.8117 |
| Net: Ut + UtT<sup>a</sup> | 0.00<br>[0.00, 0.00], <0.0010* | 0.02<br>[0.01, 0.04], <0.0010* | 0.00<br>[0.00, 0.00], <0.0010* | 1.43<br>[0.37, 5.26], 0.5938 | - | - |
| Net: Ut + UtL<sup>a</sup> | - | 0.02<br>[0.01, 0.05], <0.0010* | - | 1.25<br>[0.34, 4.44], 0.7307 | - | - |
| 1h KD: Yes (Ref: No) | - | - | - | 1.16<br>[1.11, 1.22], <0.0010* | - | 1.16<br>[1.11, 1.22],<br><0.0010* |
| Treated surface contact duration | - | - | - | 1.00<br>[1.00, 1.00], 0.0230* | 1.00<br>[1.00, 1.00], 0.0577<sup>b</sup> | 1.00<br>[1.00, 1.00], 0.0445* |

\* = significant at 5% significance level, <sup>a</sup> = net reference P3, <sup>b</sup> = borderline significance, ref = reference

**Table S2.** Number and duration of net contacts results by Tiassale mosquitoes in tests with all BBnet variants based on a linear mixed effects model. Where 95% confidence intervals and *p-values* were adjusted by Dunnett’s multiple comparison test, * = significant at 5% significance level, ^a^= net reference P3, ^b^= borderline significance, ref = reference **and** ** = at the average net number of contacts per square cm i.e. 0.07 (net*number of contacts interaction significant: *F-value* = 4.17, df = 6, *p* = 0.0045).at the Permanet 3.

| Term | Mean difference [95% Confidence Interval], p-value |  |  |  |  |  |
| --- | --- | --- | --- | --- | --- | --- |
|  | Overall |  | Overall treated surface |  | Treated surface per square cm |  |
|  | Contact duration | Number of contacts | Contact duration | Number of contacts | Contact duration ** | Number of contacts |
| (Intercept) | -652.26<br>[-1552.14, 247.63], 0.1509 | 4847.47<br>[-664.50, 10359.44], 0.0831 | -281.55<br>[-1129.41, 566.32], 0.5037 | 6304.83<br>[3558.85, 9050.81], <0.0010* | - | 0.09<br>[0.05, 0.13], <0.0010* |
| Net: P3 + P3T<sup>a</sup> | 976.45<br>[-221.15, 2174.05], 0.1073 | -3978.16<br>[-10552.70, 2596.37], 0.2284 | 845.11<br>[27.24, 1662.98], 0.0433* | -2870.22<br>[-6079.37, 338.94], 0.0778 | 0.01<br>[-0.03, 0.05], 0.9666 | -0.05<br>[-0.10, 0.00], 0.0470* |
| Net: P3 + P3L<sup>a</sup> | 1888.65<br>[750.43, 3026.86], 0.0017* | -933.98<br>[-6411.52, 4543.57], 0.7281 | 1794.62<br>[1028.98, 2560.26], <0.0010* | -1858.07<br>[-4923.00, 1206.87], 0.2259 | 0.02<br>[-0.02, 0.06], 0.6438 | -0.04<br>[-0.09, 0.00], 0.0695 |
| Net: P3 + UtT<sup>a</sup> | 908.71<br>[-293.69, 2111.11], 0.1346 | -3043.99<br>[-9759.12, 3671.13], 0.3651 | 481.38<br>[-364.03, 1326.80], 0.2547 | -3545.48<br>[-6768.82, -322.14], 0.0321* | 0.00<br>[-0.04, 0.04], 0.9995 | -0.05<br>[-0.10, 0.00], 0.0417* |
| Net: P3 + Utl<sup>a</sup> | 2646.89<br>[1504.83, 3788.95], <0.0010* | -2225.06<br>[-8203.91, 3753.80], 0.4523 | 1691.48<br>[885.42, 2497.54], <0.0010* | -4504.17<br>[-7667.04, -1341.31], 0.0067* | 0.02<br>[-0.02, 0.06], 0.5023 | -0.07<br>[-0.12, -0.02], 0.0068* |
| Net: Ut + P3T<sup>a</sup> | 1106.41<br>[-33.58, 2246.40], 0.0568<sup>b</sup> | -2786.16<br>[-8979.44, 3407.13], 0.3683 | 728.46<br>[-257.68, 1714.60], 0.1422 | -7722.39<br>[-10783.45, -4661.34], <0.0010* | 0.08<br>[0.01, 0.14], 0.0119* | -0.09<br>[-0.13, -0.04], <0.0010* |
| Net: Ut + P3L<sup>a</sup> | 3023.08<br>[1889.50, 4156.66], <0.0010* | 949.42<br>[-5033.63, 6932.48], 0.7488 | 1465.99<br>[533.44, 2398.54], 0.0031* | -6959.89<br>[-10024.82, -3894.95], <0.0010* | 0.06<br>[0.02, 0.11], 0.0012* | -0.04<br>[-0.09, 0.01], 0.0986 |
| Net: Ut + UtT<sup>a</sup> | 3021.72<br>[1778.65, 4264.80], <0.0010* | 10282.26<br>[4247.78, 16316.74], 0.0015* | - | - | - | - |
| Net: Ut + Utl<sup>a</sup> | 981.42<br>[-489.39, 2452.23], 0.1852 | 20357.59<br>[13549.12, 27166.05], <0.0010* | - | - | - | - |
| Number of contacts | 0.18<br>[0.13, 0.23], <0.0010* | - | 0.13<br>[0.05, 0.22], 0.0024* | - | - | - |
| Head: Right (ref: Left) | - | -2811.02<br>[-5729.85, 107.81], 0.0585<sup>b</sup> | - | - | - | - |
| Max no.of mosquitoes | - | 1247.64<br>[354.39, 2140.88], 0.0076* | - | 551.00<br>[17.01, 1084.99], 0.0435* | - | 0.01<br>[0.00, 0.02], 0.0212* |

\* = significant at 5% significance level, <sup>a</sup> = net reference P3, <sup>b</sup> = borderline significance, ref = reference, \*\* = at the average net number of contacts per square cm i.e. 0.07 (net\*number of contacts interaction significant: *F-value* = 4.17, *df* = 6, *p* = 0.0045).

## References

1. Churcher, T., Lissenden, N., Griffin, J. T., Worrall, E. & Ranson, H. The impact of pyrethroid resistance on the efficacy and effectiveness of bednets for malaria control in Africa. eLife, 116, https://doi.org/10.7554/eLife.16090 (2016)

2. WHO Vector control products, Pre-Qualification reports https://extranet.who.int/pqweb/prequalification-reports

3. Gleave, K, Lissenden, N., Chaplin, M., Choi, L. & Ranson, H. Piperonyl butoxide (PBO) combined with pyrethroids in insecticide-treated nets to prevent malaria in Africa. Cochrane Database Syst Rev 24, 10.1002/14651858.CD012776.pub3

4. Dagg, K., et al. Evaluation of toxicity of clothianidin (neonicotinoid) and chlorfenapyr (pyrrole) insecticides and cross-resistance to other public health insecticides in Anopheles arabiensis from Ethiopia. Malaria J. 18, 1849 https://doi.org/10.1186/s12936-019-2685-2

5. Anopheles gambiae 1000 Genomes Consortium. Genome variation and population structure among 1142 mosquitoes of the African malaria vector species Anopheles gambiae and Anopheles coluzzii. Genome Research 30, 1533–1546. 10.1101/gr.262790.120. (2020)

6. Coleman, M., et al. Developing global maps of insecticide resistance risk to improve vector control. Malaria J. 16, pp.1–9. (2017)

7. Murray, G. P. D., et al. Barrier bednets target malaria vectors and expand the range of usable insecticides. Nat Micro, 5, 40–47 https://rdcu.be/bZ6ip (2019)

8. Parker, J. E. A., et al Infrared video tracking of Anopheles gambiae at insecticide-treated bed nets reveals rapid decisive impact after brief localised net contact. Sci. Reps, 5, 13392. Do10.1038/srep13392 (2015).

9. Angarita-Jaimes, N. et al. A novel video-tracking system to record behaviour of nocturnal mosquitoes attacking human hosts. J. Roy Soc Interface 13, https://dx.doi.org/10.1098/rsif.2015.0974 (2016).

10. Sutcliffe, S. & Yin, S. Effects of indoor air movement and ambient temperature on mosquito (Anopheles gambiae) behaviour around bed nets: implications for malaria prevention initiatives. Malaria. 427. https://doi.org/10.1186/s12936-021-03957-y (2021).

11. Parker, J. E. A., et al. Host-seeking activity of a Tanzanian population of Anopheles arabiensis at an insecticide treated bed net. Malaria J. 16, DOI 10.1186/s12936-017-1909-6 (2017).

12. Sherrard-Smith, E., et al. Mosquito feeding behaviour and how it influences malaria transmission across Africa PNAS USA 116, 15086 https://doi.org/10.1073/pnas.1820646116 (2019).

13. Firth, D. Bias reduction of maximum likelihood estimates. Biometrika, 80, pp.27–38 (1993)

14. R Core Team. R: A language and environment for statistical computing. R Foundation for Statistical Computing, Vienna, Austria. URL https://www.R-project.org/. (2022)

15. Posit team. RStudio: Integrated Development Environment for R. Posit So∼ware, PBC, Boston, MA. URL http://www.posit.co/. (2023)

16. Bates, D. et al. Fitting Linear Mixed-Effects Models Using lme4. Journal of Statistical Software, 67, 1–48. doi:10.18637/jss.v067.i01 (2015)

17. Lenth, R. emmeans: Estimated Marginal Means, aka Least-Squares Means_. R package version 1.8.4-1, < https://CRAN.R-project.org/package=emmeans>.(2023)

18. Hartig, F. DHARMa: Residual DiagnosMcs for Hierarchical (MulM-Level / Mixed) Regression Models_. R package version 0.4.6, https://CRAN.R-project.org/package=DHARMa (2022)

19. Wickham, H. ggplot2: Elegant Graphics for Data Analysis. Springer-Verlag New York, (2016)

20. Tang Y, Horikoshi, M, and Li, W. “ggfortify: Unified Interface to Visualize Statistical Result of Popular R Packages.” The R Journal 8.2 :478–489 (2016).

